# Cuttlefish interact with multimodal “arm wave sign” displays

**DOI:** 10.1101/2025.04.13.648584

**Authors:** Sophie Cohen-Bodénès, Peter Neri

## Abstract

In addition to the well-known extraordinary changes in visual appearance they can generate at the level of their mantle, cuttle-fish can produce various body configurations combining chromatic, postural, and locomotion patterns, both for camouflaging and communication. We introduce a previously undescribed communication display in two cuttlefish species: Sepia officinalis and Sepia bandensis. The four “arm wave signs” are stereotyped arm movements consisting of long-lasting, expressive, and repeated sequences of undulations of the arms, which can be combined and expressed following specific patterns. Using non-invasive behavioral experiments, we tested the hypothesis that they represent multimodal communication displays. To assess the role of visual cues, we recorded videos of animals signing and played them back to individual cuttlefish participants. When seeing the movies, cuttlefish waved back at the display. Most importantly, they were more likely to wave back when the movie was in upright (original) configuration as opposed to flipped upside-down, similar to the manner in which humans perceive faces and other socially relevant displays. In addition to their visually striking display, arm wave signs produce mechanical waves in the water, prompting us to explore the possibility that they may also be perceived via mechanoreception. Using playback experiments similar to those adopted in vision, we obtained preliminary evidence to support this hypothesis, indicating that arm wave signs may represent multimodal signals involving vision and mechanoreception. Our new result on communication with arm wave signs opens up novel possibilities for understanding vibration-mediated communication through the lateral line and/or the statocysts in a cephalopod species as an example of convergent evolution with vertebrates.

## Introduction

The cryptic abilities of cuttlefish, known as “chameleons of the sea”, are justly celebrated as stunning examples of animal camouflage. For example, Sepia officinalis can combine more than fifty body patterns to approximate specific features of artificial backgrounds (1–6). Skin patterning is a versatile ability that is also used for communication (7), the intense zebra bands being the most representative pattern during reproductive behaviors in Sepia officinalis (8). Cuttlefish can also use a variety of body and arm postures to communicate information about their intention to fight or mate (8). For example, the “raised arms” are associated with the “flamboyant display” to intimidate predators, while the “extended fourth arm” is used by male cuttlefish to communicate their intention to mate (7–9). Cuttlefish also alter the shape of their body: they can flatten their body to appear larger (9) and frighten a potential predator, as is the case in the deimatic display (7, 9, 10). More generally, arms can be “raised, lowered, splayed, split, V-curled, or contorted” as summarized by Mather (7). Arms can make the animal look larger and can be used to frighten and repeal as well as look more dominant.

Several functions have been suggested for behaviors that use arms in cephalopods, notably masquerade, flamboyant displays, and luring (7, 8). In cuttlefish and octopus, specific arm movements and postures have also been described and interpreted as mimicking behaviors. In cuttlefish, Okamoto and collaborators identified and studied a unique arm-flapping behavior in Sepia pharaonis to which they attributed the function of crab mimicry, especially during hunting (11). However, this study did not test the hypothesis that this remarkable display could also be addressed from one cuttlefish to another for communication. Norman had previously shown that arm postures and movements are used by octopus to mimic the behavior of sole, sea snake, and lionfish (12). However, as noted by the authors, there were no behavioral studies that complemented those findings to test the hypothesis of mimicry. In squids, communication through arm postures/movements and dances/choregraphies have been extensively studied and documented, notably in the form of ethograms (7, 13–15).

Cephalopods in general communicate multimodally by combining various sensory modalities: visually with body patterns (5, 16–21), via tactile stimulation (7, 22–24), chemically with ink or pheromones (25–31), and through light polarization (32–36). Cuttlefish do not have ears, but perceive sounds and low vibrations through their lateral line hair cells and their statocysts (10, 37–47). There is currently no clear evidence pertaining to the possibility that cuttlefish may interact with each other through sounds (to be understood as low-frequency vibrations) as vertebrate teleost fish or cetaceans do (7, 48). However, it is established that they respond behaviorally to sounds ranging from 20 to 1000 Hz (38, 40–47, 49–52). The classical behaviors reported with sound stimulation are: startle response (53), inking, jetting, and rapid coloration change, moving, burrowing, orienting, swimming, and body patterns associated with specific frequencies such as dark flashing, bleaching, and deimatic display (40, 41, 43, 44, 51, 54). In squids, several escape responses (inking, jetting, fleeing, deimatic display) evoking predator evasion have been revealed (55). As Mather stated, despite the lack of evidence so far, it is possible that cephalopods can detect vibrations created by conspecifics using the lateral line to facilitate social behavior, shoaling at night (7). To our knowledge, there is currently no evidence of communication through arm movements that would be perceived by mechanoreception (sound/vibration) in cephalopods.

In amphibians and fish, the lateral line is involved in prey detection, predator avoidance, rheotaxis (56), the localization of stationary objects, and schooling behavior (57, 58). It also provides a sense of spatial awareness for the animal and participates in its ability to navigate the surrounding environment. Evidence from comparative ecology in fish and amphibians, which are the two other groups equipped with a lateral line organ, supports the notion that the lateral line plays a role in communication (59–63). The cuttlefish lateral line and statocysts are an example of convergent evolution between an invertebrate and a vertebrate sensory system (37, 45). In cuttlefish, the attested function of the lateral line is the detection of small movements of water conveyed by hydrodynamic pressure gradients (37, 39, 64, 65). Cephalopods can perceive long-range sounds through their statocysts, particle-motion detectors (43) akin to the cochlea in vertebrates (41, 42, 45– 47, 50). In cephalopods specifically, the function of the lateral line and the statocysts has been established for navigation, detection of prey, predator avoidance, and, hypothetically, social schooling behavior (39, 66, 67).

In this study, motivated by the evidence reviewed above, we report a previously undescribed category of arm wave displays in two species of cuttlefish (Sepia officinalis and Sepia bandensis), which we refer to as “arm wave signs” (**Figure** 1, **Supplementary Videos 1–4**). We hypothesized that those signals are communication displays that cuttlefish use to interact with each other due to their specific features. The four “arm wave signs are stereotyped arm movements consisting of repeated, expressive, long-lasting sequences of undulations of the arms, which can be combined and expressed in correlation with other body patterns, including classically described dynamic skin body coloration for communication.

After presenting specific ethograms of the signs in two cuttle-fish species, we present the results of two non-invasive behavioral experiments. First, we tested the hypothesis that these arm movements are informational displays used by cuttlefish to interact with one another visually. Our setup consisted in randomly presenting to the cuttlefish videos of themselves signing in upward (normal) and upside-down (control) configurations.

Prompted by the results obtained in the visual domain, we tested the hypothesis that arm waves may be perceived via mechanoreception, given that they produce mechanical waves in the surrounding water. The lateral line and the statocysts are the two candidate organs in cuttlefish for the perception of low-frequency movements generated by the arm wave signs (37, 39). We recorded traces of arm wave displays with a hydrophone (for the common cuttlefish) and a speaker (for juvenile Sepia bandensis) and played the traces back to the animal in normal configuration as well as two control configurations (reversed and scrambled). The play-back method has been widely used in ethological studies as a way to project back a sound of communication to an animal and record their behavioral responses (68, 69). Several studies have investigated the behavioral responses of cephalopods to sound projected by underwater speakers, and have shown that statocysts perceive particle acceleration (40– 42, 47, 51, 54, 70). To our knowledge, our study is the first to introduce hydrophone/speaker-mediated playback as a method to induce specific arm movements in cuttlefish.

Below we first present results from the non-invasive behavioral experiments introduced above, which provide the first evidence that arm wave signs may represent communication signals perceived visually and through mechanoreception. In the discussion section, we address the implications of our study for the study of communication through arm movements in cephalopods. We finally open the debate on the potential role of the lateral line and statocysts in perceiving arm wave signs in cuttlefish, and the possible function of mechanoreceptors in communication behavior as an example of convergent evolution between cephalopods and vertebrates.

## Methods

### A. Animals and husbandry

We conducted two types of behavioral experiments: “visual” and “mechanosensory”. For the visual experiments, we tested eight adult cuttlefish Sepia officinalis (3–12 months) and ten juvenile cuttlefish Sepia bandensis (1–2 months). For the mechanosensory experiments, we tested eight adult Sepia officinalis cuttlefish and eight juvenile Sepia bandensis cuttlefish. All animals were reared from eggs and kept in hatchlings following established guidelines (71), at the authorized institutional supplier Aquarium de la Rochelle (La Rochelle, France). The eggs were collected in the Atlantic Ocean (for Sepia officinalis) or in the Indo-Pacific Ocean (for Sepia bandensis). The cuttlefish were reared in captivity (hatched from the eggs) by a team of biologists specialized in the breeding of cuttle-fish. Transportation was optimized by oxygenation enrichment, noise and vibration minimization in a cool specialized container without light exposure, and following food deprivation prior to transport to avoid accumulation of toxic ammonia level. Once the eggs / juveniles reached our facility, they were acclimatized in four artificial marine water tanks (300 L each) for 2 hours. When handling eggs, we introduced additional air pumps to provide sufficient oxygenation for their development.

We ensured that the tank environment approximated the natural habitat as closely as possible, while at the same time providing adequate monitoring of water quality and animal health and well being. The tanks operated a closed-water system that allowed constant monitoring and optimization of water parameters and contained various oxygen pumps, a skimmer, mechanical and biological filtration, and carbon for ink removal. We introduced sand enrichment (7-cm layer of natural marine uniform sand with granulometry of 1–4 mm, enriched with natural pebbles of different size/color placed randomly across the tank), and live rocks (*∼* 15 live rocks of different sizes for a total of *∼* 20 kg per tank) to provide shelter, and more generally to encourage cuttlefish expression of their full behavioral repertoire (burying in sand, establishing territories and dominance hierarchies, protection from lighting within areas shadowed by rocks, swimming by jet propulsion).

The tanks were visually isolated from each other to prevent cuttlefish from interacting with animals in other tanks, and tank surface was opacified with adhesive vinyl sheets to minimize mirror reflections. We established a light/dark cycle of 12-hours/12-hours, and only conducted experiments during the day. Live food was provided three times a day: at 9 a.m., at 2 p.m., and at 8 p.m. We never deprived animals; instead, food was systematically provided so that cuttlefish could hunt at will (especially at night) and would not engage in aggressive behavior for food competition. From hatchlings to 1 month of age, cuttlefish were fed a mixture of live Artemias and live Mysids. Once they reached 1 month of age (2 month for Sepia bandensis), they were fed larger live grass marine shrimp (*Palaemonetes vulgaris*) and live crabs.

### B. Ethical Statement

In this study, we only performed non-invasive observational experiments (never involving any invasive procedure, pain, or lasting harm to the animal). In compliance with 3Rs principles (72) and with established guidelines for the use and care of cephalopods in captivity (71), we housed animals in large tanks enriched with sand, rocks, sheltered areas, and provided only live food at will to allow expression of their full behavioral repertoire, notably hunting at will (Refine). We allowed a maximum of six animals per tank. We managed to establish a trade-off between sufficient statistical power and reduction of the number of animals used per experiment (Reduce). Our observational non-invasive experiments are therefore classified below the threshold for explicit ethics approval as expressed in Directive 63/10. However, we sought and obtained validation from the head of the ethics committee of our Biology Department (IBENS, École Normale Supérieure, Paris, France).

### C. Experimental procedure

#### C.1. Visual experiments: stimuli and presentation protocol

Our first goal was to record sequences of spontaneous expression of the four arm wave signs in a variety of behavioral contexts. When a cuttlefish settled or moved, signing, within a specific area of its choice, we placed an underwater camera (GoPro Hero13) above the tank to record mantle activity for *∼* 2 hours at a time, three times a day, over a period of *∼* 9 months. To obtain frontal video footage of the face and arms, we placed an iPad Pro Retina (iPad Pro Retina 12.9) in front of the tank to record cuttlefish waving from a standardized point of view. To construct our stimuli, we selected sequences of signs from each individual over several (1–7) months filmed from various viewpoints, spanning a wide range of body patterns (See **Supplementary videos S5**). Starting from this large dataset, we then randomly selected sequences during which cuttlefish elicited specific wave signs, separated by sequences during which they did not sign. For the Sepia officinalis cohort, sequences were displayed against the black background of a MacBook Pro 13” screen (Intel Iris, 2GHz Intel Core-i running on MacOS Sequoia 15.3.2), with the stimulus occupying half the available screen. We generated 18 sequences displaying 13 waving animals (6 males and 7 females, aged 3–7 months), separated by presentations of cuttlefish not signing. The assembled video for final presentation contained 61 signs and lasted 10 minutes. We randomly presented the normal “upright” version of the video and a control “upside-down” version, in which the video was flipped upside-down.

For the Sepia bandensis cohort, we selected four versions of arm wave signs (two versions of the “up” sign and two versions of the “crown” sign). In addition to the upside-down version of the videos, we created two other control categories: non-signing (two recordings of cuttlefish standing still, two recordings of cuttlefish swimming), and non-signing upside-down (see **Supplementary videos S7**). We displayed an equal number of exemplars from each of the four categories (“signing”, “signing upside-down”, “no signing”, “no signing upside-down”) in random sequence for 20 minutes, against the black background of a MacBook Pro 13” screen (stimulus occupied 60% of screen). Each exemplar lasted 5 seconds, and exemplars were separated by a pause video displaying the empty tank.

We ran 5–10 trials a day (one trial meaning one 20-minute sequence as detailed above) over a period of 1 month. On average, we tested three randomly selected animals per day. Animals were placed in the testing arena tanks for sessions lasting two hours maximum, to prevent both animal fatigue and habituation. Before testing, we allowed each animal to acclimatize to the test tank for 30 minutes, to prevent any stress-related bias. We did not introduce conditioning or reinforcement (e.g reward). We allowed breaks between trials to allow the animal to rest and prevent early habituation to stimulation.

#### C.2. Mechanosensory experiments

##### Stimulus design/creation

We recorded sequences of spontaneous expression of wave signs from one cuttlefish Sepia officinalis (eight months) and one juvenile Sepia bandensis (1 month), which we then played back to our test animals. To record the vibrations emitted by the arm waves, we used a hydrophone Brüel and Kjaer Type 8103 for Sepia officinalis and Aquarian Audio hydrophone model AS1 Scientific for Sepia Bandensis. We isolated seven instances of the “up” wave sign from the first individual, which we cropped to *∼* 10 seconds each (maximum sign duration in Sepia officinalis), and eight instances of the “up” and “crown” signs from the second individual, which we cropped to *∼* 5 seconds each (maximum sign duration in Sepia bandensis).

For each recorded waveform, we generated two control waveforms: reversed and scrambled. The “reversed” waveform was obtained by simply time-reversing the original trace. This manipulation is relevant because it can lead to a loss of interpretability for meaningful sounds, while largely preserving the low-level content of the original trace. For example, speech becomes unintelligible when played backwards (73, 74). To obtain the “scrambled” waveform, we computed the Fourier transform of the original waveform, replaced its phase spectrum with one obtained from a random trace in which each time point was randomly assigned a value from a Gaussian distribution, and then computed the inverse Fourier transform to obtain the corresponding waveform. In this manner, the scrambled waveform retains the same power spectrum associated with the original waveform, but loses the original phase structure. This manipulation is interesting because it removes important features of natural signals, while preserving the amount of overall intensity associated with the original stimulus. For example, interference with the power spectrum of natural scenes only causes small degradation of recognizable content, provided the phase spectrum is left unaltered. In contrast, interference with the phase spectrum renders the content of natural scenes uninterpretable, even though the power spectrum was left unaltered (75). We then generated pseudo-random sequences of the three stimuli detailed above: an initial pause of 10 seconds, followed by a stimulus, another 10-second pause, another stimulus, and so on, for a total of 21 stimuli over a period of approximately 7 minutes (430 seconds). Each stimulus was randomly selected to be the normal waveform, the backward waveform, or the scrambled waveform in such a way that each waveform type was presented seven times during the 7-minute sequence.

##### Presentation protocol

We played back the above-detailed stimuli to 8 Sepia officinalis participants via a hydrophone, and to 10 Sepia bandensis participants via an underwater speaker equipped with sub-woofers.

When testing Sepia officinalis, we did not interfere with their daily routine. We let animals behave naturally in their home tank to encourage expression of natural and spontaneous behavior. We waited for a cuttlefish to settle in a specific area of its choice, and tested the animal when the hydrophone (Brüel and Kjaer Type 8103) was at a distance of approximately 3–10 cm from its arms (one body length for adult cuttlefish Sepia officinalis). An underwater camera (GoPro Hero7) was fixed in advance above the tank to record mantle activity and dynamic skin patterns.

When testing Sepia bandensis, we isolated the animal within an ad-hoc test tank. We randomly picked a juvenile cuttle-fish from the home tank and let the animal acclimate to the new tank for 30 minutes prior to testing (same protocol as for visual experiments). The speaker (Ortizan X8 Pro or TOZO PA1), enclosed in a thin waterproof bag for water protection, was placed in the test tank before introducing the animal.

We played back stimuli in pseudo-random sequence for both hydrophone-based and speaker-based experiments. The displayed sequence was recorded for subsequent blind-alignment with behavioral scoring. We placed an iPad Retina Pro in front of, and a camera GoPro Hero13 above, the test animal, to record its behavioral responses to the mechanosensory stimuli.

#### C.3. Data Analysis

##### Visual experiments

For Sepia officinalis, we collected 128 trial-associated video recordings from 8 cuttlefish (aged 8–12 months), who carried out 10–15 trials per day for a period of 2 months. For Sepia bandensis, we collected 52 trials from 10 animals (aged 1 month), who carried out 10–15 trials per day for a period of 1 month.

Each video recording of the animal behaving in response to the visual stimuli was scored manually, by annotating the time of production and identify (which of the four identified types) each produced wave sign. This information was recorded without knowledge of the video sequence presented to the animal during the video-recorded trial, i.e. labeling was performed blindly. A custom script loaded the labeled file detailed above, and automatically matched it against a separate file recording timestamps for the visually displayed signs. This procedure was adopted to ensure that stimulus presentation and behavioral scoring were carried out independently, and only matched automatically by computer software.

##### Mechanosensory experiments

For Sepia officinalis, we collected 113 videos associated with the trials from 8 cuttlefish (aged 8–12 months), who carried out 10–15 trials per day for a period of 3 weeks. For Sepia bandensis, we collected 49 trials from 10 animals (aged 1 month), who carried out 10–15 trials per day for a period of 2 weeks.

Each video recording of the animal behaving in response to mechanosensory stimuli was scored manually using the same procedure adopted for the visual experiments (see above), and similarly matched against the previously recorded stimulation sequence via an automated computer algorithm.

For Sepia officinalis, we also collected mantle activity via the GoPro camera. We manually annotated the expression of strong highly contrasted skin body coloration patterns. We annotated as communication displays those instances in which the color/brightness of the body pattern changed vividly and rapidly. For example, we would label a time window in which the cuttlefish would exhibit a bright orange pattern or a vivid black point in response to playback stimuli. We took as a reference the coloration classically referenced as skin body coloration displays in the literature on cuttlefish Sepia officinalis (2–9, 21, 76).

##### Statistical analysis

For both experiments, we carried out ANOVAs, t-tests, Wilcoxon tests, and Friedman tests.

## Results

### Ethograms of the four arm wave signs

Extensive observations over several months on two successive colonies of 24 cuttlefish Sepia officinalis led to the identification of four stereotyped “arm wave signs” displayed in different behavioral contexts. These displays involve long-lasting (2– 10 seconds) movements of the arms, exhibited in repetitive sequences where different signs can be combined with other signs. We refer to the four different signs with the terms “up”, “side”, “roll”, and “crown” (**Figure 1**). They are often combined with the expression of various chromatic body patterns and locomotor components, such as fin undulation/contraction (**Figure 1A**). They can be expressed by both juveniles and adults, males and females.

**Fig. 1.**
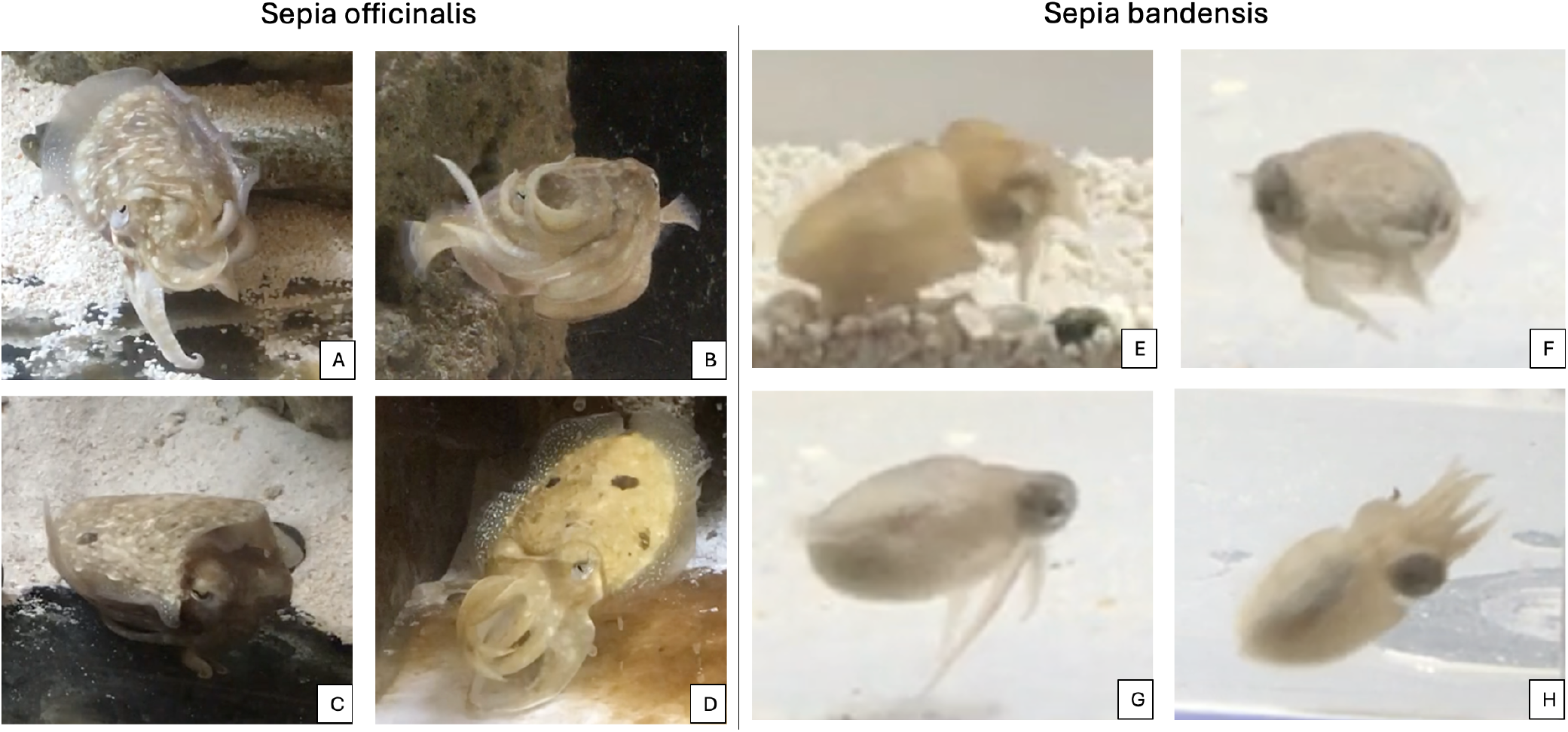
The four arm wave signs in adult Sepia Officinalis (A–D) and juvenile Sepia bansensis. **(E–H)** The “up” sign (**A**,**E**), most frequently expressed, involves extension of the first pair of arms in the upward direction and extension of the fourth arm pair, while the second and third arm pairs are twisted in the middle. When expressing the “side” sign (**B**,**F**), cuttlefish roll all arms to one or the other side of the body. In the “roll” sign (**C**,**G**), all arms are rolled under the head to change its shape and emphasize the eyes. Finally, the “crown” sign (**D**,**H**) involves a “spitting” movement with the arms arranged in the shape of a crown. Cuttlefish exhibit these wave signs in different behavioral contexts such as sitting (**A**,**B**,**C**) or swimming (**B**,**G**,**H**), often in combination with high-contrast skin patterns (black points in **C**, orange coloration with black spots in **D**) and contraction of the fins (**C**). See **Supplementary Videos 1–4, 5**.

We observed that cuttlefish sometimes appear to direct these signs not only at conspecifics, but also towards prey, and occasionally humans approaching the tank. In addition to being expressed during interactions, they can be associated with various behavioral states that do not involve specific sensory stimulation, such as swimming, eating, resting, and burying. They can be expressed during hunting, and as aversive signals during the introduction of an unfamiliar object (such as a net) in the tank. We postulated and experimentally tested that these arm waves can function (among other things) as intraspecific communication signals used by cuttlefish to interact with each other. Their highly visible and exaggerated nature suggests that they are costly and unrelated to camouflage (77, 78). Because costs come with trade-offs, it is likely that they serve a useful function in terms of fitness, for example in conveying useful information during mating.

We identified variants of these signs in a different cuttlefish species, Sepia bandensis (**Figure 1B**). These animals frequently express the “up” and “crown” signs from early juvenile stage (1 week). The “crown” and “roll” signs appear to serve as defensive displays in these juvenile cuttlefish. The crown sign is often repeated several times in a sequence, and is associated with dark skin coloration.

### Arm wave signs are reciprocated in response to visual displays

We performed visual experiments in two distinct cohorts of cuttlefish: 8 adult cuttlefish Sepia officinalis, and 10 juvenile Sepia bandensis. We adopted slightly different experimental protocols for the two cohorts, because the results obtained with the first cohort informed our subsequent design of the protocol deployed to the second cohort.

For our tests with Sepia officinalis, the animals were shown visual displays of arm wave signs recorded from a conspecific (**Figure 2A**–**B**), presented in either upright or upside-down configuration (in the latter, the video was flipped upside-down). We observed that the test animal (**Figure 2B**) responded more often when the stimulus was in upright configuration (red segments in **Figure 2C**) as opposed to upside-down configuration (blue segments in **Figure 2C**).

**Fig. 2.**
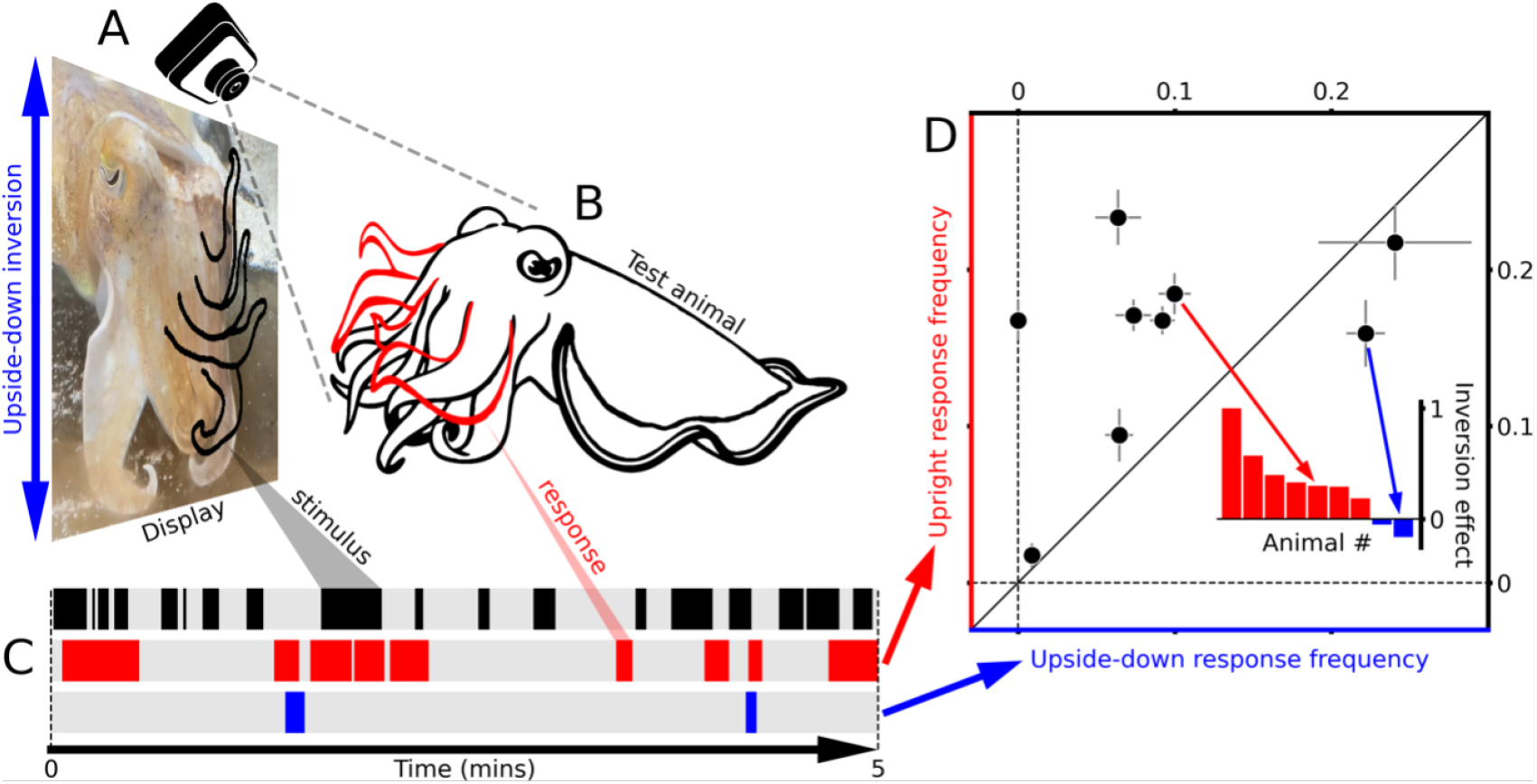
Cuttlefish Sepia officinalis wave back at movies of other cuttlefish waving. **A** Footage of cuttlefish producing wave signs was played back to a real cuttlefish **B** looking at a flat screen, while the animal was being simultaneously recorded (camera in **A**). This setup made it possible to align occurrences of wave signs in the visual stimulus (indicated by black segments in **C**) with occurrences of wave signs produced by the test animal (indicated by red segments in **C**). As indicated by the 5-minute excerpt in **C**, it appears that the test animal produced wave signs in response to those delivered by the visual stimulus, possibly with some delay (see **Figure S1**). Furthermore, when the visual stimulus was inverted upside-down during playback (blue double-headed arrow in **A**), the test animal produced fewer signs (blue segment in **C**). We quantified this effect by calculating the frequency of signing in the upright configuration, plotted on the y axis in **D**, versus the frequency of signing in the upside-down configuration, plotted on the x axis in **D**, separately for each animal (one data point per animal). Data points that fall above the diagonal unity line indicate more frequent response in the upright configuration compared with the upside-down configuration, while points that fall below the diagonal unity line indicate the opposite trend. To further quantify potential differences between upright and inverted configurations, we computed their normalized difference (see main text) in the form of an inversion effect, plotted in the inset to **D**, separately for different animals. Error bars in **D** plot ±1 SEM.

We quantified the above observation by measuring the probability of wave signing for different animals. When computing this probability, it was necessary to take into account the stimulus-response delay involved in the signing process: it is reasonable to expect that cuttlefish would not sign back at the visual stimulus immediately because, at the very least, they would need time to process such a complex stimulus and initiate a corresponding motor response on their own part. In other words, it would be unrealistic to expect that the red segments would perfectly align with the black segments in **Figure 2C**, even for noiseless measurements of a perfectly responding system. To estimate the delay involved, we performed a behavioral variant of the spike-triggered analysis used in electrophysiology to characterize the temporal impulse response of sensory neurons (see caption to **Figure S1** for details). Using this approach, we identified a sluggish response spanning approximately 15 seconds of temporal integration (positive portion of the trace in **Figure S1A**). We then incorporated this knowledge into the analysis presented in **Figure 2D**: before computing the signing frequency during stimulation epochs, we convolved the stimulus trace with a 15 second temporal integration window that effectively delayed and smeared the stimulus sequence in accordance with **Figure S1A**.

Each dot in **Figure 2D** represents an animal participant, with the y coordinate of each dot plotting the probability of signing for the corresponding animal in response to the upright display, and the x coordinate plotting the probability of signing in response to the upside-down display. Our measurements show a tendency to fall above the diagonal unity line, indicating that signing was more frequent in the upright configuration compared with the upside-down configuration. To further quantify this effect and test its statistical significance, we computed the normalized difference between upright and upside-down configurations in the form of (*y − x*)*/*(*y* + *x*), where *y* is the value on the y axis and *x* is the value on the x axis in **Figure 2D**. This index, which we term the “inversion effect”, is 1 if animals only sign for the upright configuration (*x*=0), 0 if they sign equally for both configurations (*y* = *x*), and -1 if they only sign for the inverted configuration (*y*=0). As shown in the inset to **Figure 2D**, the inversion effect shows a significant tendency towards positive values across animals (Wilcoxon test for these values to differ from 0 returns p*<*0.02).

We sought to replicate and validate the results obtained from Sepia officinalis in a different species, Sepia bandensis, and for a younger stage of development (juvenile). We observed that juvenile Sepia bandensis display arm wave signs that are comparable to those produced by Sepia officinalis (**Figure 1**). We individually tested ten Sepia bandensis participants, each placed inside a test tank in front of a visual display. The animals showed a similar tendency to sign more frequently in response to the upright display compared with the upside-down display. For these experiments, we also included two additional stimulus configurations: cuttlefish not signing in upright configuration, and cuttlefish not signing in upside-down configuration. Test animals rarely signed in response to these configurations (**Figure 3**), further demonstrating the specificity of the signing response. These effects were confirmed by a repeated measured one-way ANOVA against the 4 conditions (p=0.0007), corroborated by a two-tailed paired t-test between conditions upright and upside-down (p=0.0147), upright and no signing upright (p=0.0040), upright and no signing upside-down (p=0.0036). No significant effect was found for paired comparisons involving the three configurations of upside-down, no signing upright, and no signing upside-down.

**Fig. 3.**
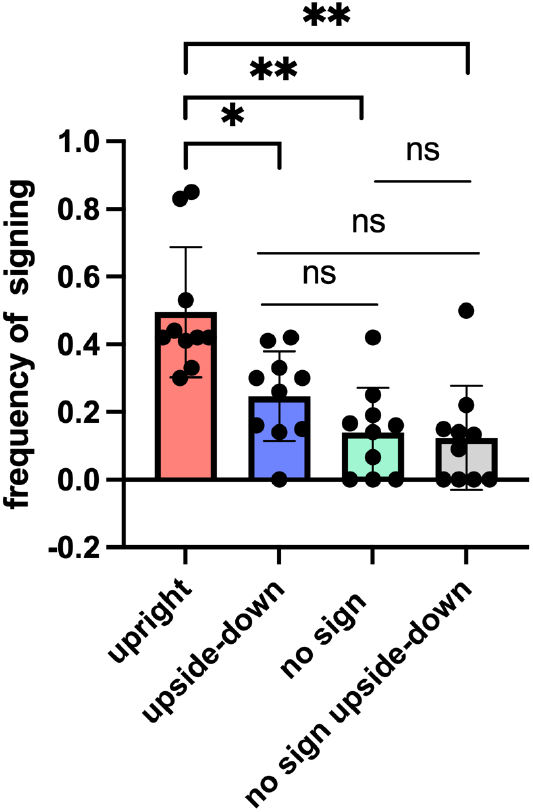
Sepia bandensis wave back at movies of themselves waving “up” and “crown”. Each dot refers to a different animal. Bars plot the frequency of signing back at the visual display for the four different stimulus conditions tested in the visual experiments (x axis, see Methods). Significance of differences between conditions is indicated by *∗* (p*<*p=0.02), *∗∗* (p*<*0.004), ns (not significant).

### Cuttlefish wave back at mechanosensory playback of arm wave signs

Using a hydrophone, we recorded mechanosensory traces of the “up” sign from Sepia officinalis, and “up” and “crown” signs from Sepia bandensis. We then played back filtered versions of these traces via a hydrophone when testing Sepia officinalis, and via a speaker when testing Sepia bandensis. Our hypothesis was that, because signs create mechanical waves in the water, they would be perceived via mechanoreception for communication. In this section, we present results supporting this notion: the low-frequency vibrations associated with traces of the wave signs elicit the expected wave sign response in both species of cuttlefish.

We studied the probability of signing for each animal in response to three waveform types: normal, reversed, and scrambled. The “normal” configuration involves playing back the original trace (filtered to exclude high-pass noise). The “reversed” configuration involved playing the same trace in time-reversed fashion (like playing a song backward). The “scrambled” configuration involved removing all phase structure from the original trace, while preserving the same power across frequencies. This manipulation results in loss of interpretability for meaningful signals such as images and sounds, while retaining the same level of contrast/intensity.

**Figure 4** illustrates our experimental setup. **Figure 4I** shows a 3-minute sequence during which the three types of stimuli are played in pseudo-random order. **Figure 4J** shows example responses in one animal: magenta segments indicate instances of sign production on the part of the test animal in response to the stimulus sequence in **Figure 4J**.

By comparing stimulus and response sequences, we can measure the probability of signing for the different types of stimulus events.

**Fig. 4.**
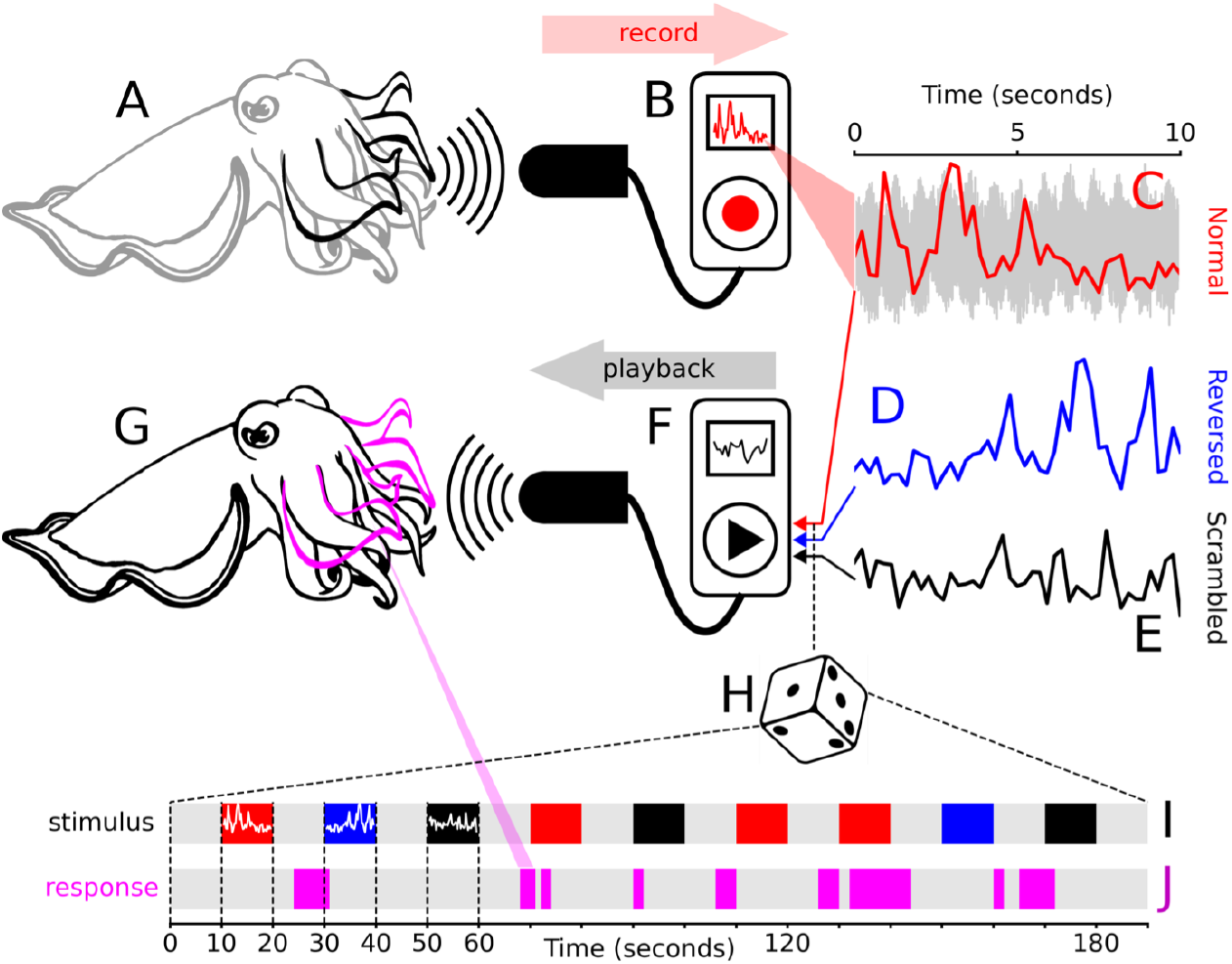
Experimental design for studying wave sign production in response to mechanosensory stimulation with a playback display. To create our stimuli, we recorded the pressure waves created by cuttlefish **A** by placing a hydrophone **B** in the vicinity of their tentacles during wave signing. To visualize relevant structure contained within the recorded trace, we have superimposed onto the original recording (gray trace in **C**) a plot of power within the range 10–30 Hz (red trace in **C**; see also **Figure S2**. For each sample of the wave sign, we created two additional stimuli: a reversed stimulus **D**, obtained by simply time-reversing the original trace, and a scrambled stimulus **E**, obtained by randomly scrambling the phase spectrum of the original trace while preserving its power spectrum (see main text). We then played back these stimuli in pseudo-random order (**H**–**I**) to a test animal in the tank (**F**–**G**), and recorded wave sign responses from the test animal using the same procedures adopted in **A**. During stimulation, each stimulus (indicated by colored segment in **I**) lasted 10 seconds, followed by a 10-second pause (indicated by gray segments in **I**), and then the next stimulus. Color coding in **I** corresponds to **C**–**E**: red for the original (normal) trace, blue for reversed, and black for scrambled. We video-recorded the test animal to identify instances of sign production (magenta in **G**–**J**), which could then be matched against the stimulus sequence **I**–**J**.

**Figure 5** plots the resulting probabilities of signing for the three different types of stimulus. Animals sign more in response to the normal waveform when compared with reversed/scrambled waveforms. To confirm the significance of this data pattern, we applied a Friedman test to determine whether there was any difference in signing frequency across the three conditions, and obtained p*<* 0.003. We then proceeded with a Conover post-hoc test to evaluate pairwise differences between conditions (corrected for multiple comparisons), which returned significant differences (p*<* 0.01) for normal-versus-scrambled and normal-vs-reversed, but not difference (p= 0.35) for scrambled-versus-reversed.

**Fig. 5.**
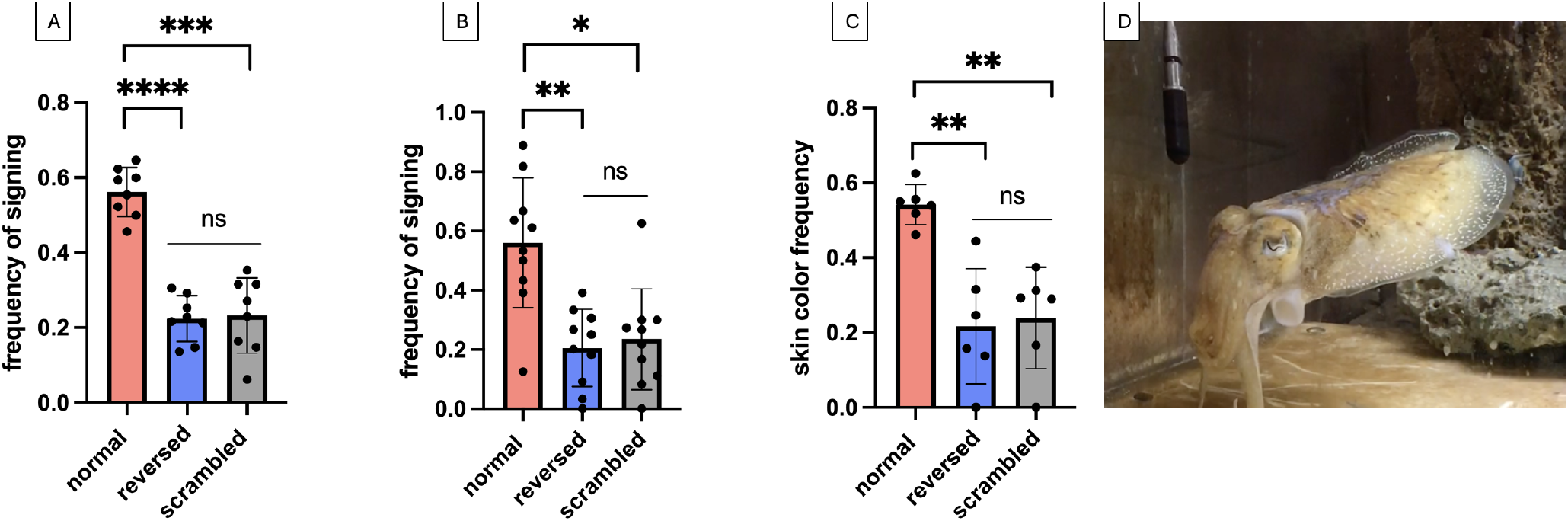
Mechanosensory playback of wave signs selectively elicits wave sign responses. **A**/**B** plot frequency of signing back in Sepia officinalis/Sepia Bandensis for the three different stimulus conditions tested in the mechanosensory experiments. **C** plots frequency of body coloration changes in Sepia officinalis (an example is shown in **D**). Plotted to the conventions of **Figure 3**.

We obtained similar results for a different species, Sepia bandensis, using an underwater speaker instead of a hydrophone for playback stimulation (for these experiments, we randomly selected the “up” or “crown” waveform for each stimulus presentation). The majority of the animals (9 out of 10) showed a significant tendency to sign more frequently in response to the normal playback stimulation when compared with reversed and scrambled conditions (repeated measured one-way ANOVA against the three conditions returns p=0.0072, corroborated by a two-tailed paired t-test between condition normal versus reversed at p=0.0064, normal versus scrambled at p=0.021, with no significant effect for reversed versus scrambled at p=0.67; see **Figure** 5), confirming the results obtained in Sepia officinalis.

Finally, we imaged skin patterns using a GoPro camera fixed above the tank. We manually annotated the expression of communication skin patterns following the same method adopted for arm signing. More specifically, we annotated communication displays in which the color/brightness of the body pattern changed vividly and rapidly. For instance, we would label a time window where the cuttlefish would exhibit bright orange pattern or vivid black point, or a traveling wave in response to the playback stimuli.

We found that all cuttlefish expressed significantly more correlated high-contrast skin patterns during the “normal” condition (**Figure** 5 **C-D**); see also **Supplementary video 5)**. We also observed that, in addition to expressing vivid skin patterns, Sepia officinalis displayed specific behaviors and body patterns during the playback experiment such as lateral pupil contraction, eye and head movements, burying, swimming toward the hydrophone to touch the device, extending the tentacules as in a prey-catching attempt displaying “hunting arms”, extension of the fourth arm associated with Zebra bands, change in body shape, and inking. We also observed recurrence of fin flipping in a stereotyped manner similar to the wave signs. In Sepia bandensis, we observed rapid changes in coloration (usually black), hunting arms (consisting of waving the first pair of arms on one side or the other), inking, swimming, body contraction, especially of the mantle, eyes and head movements.

### Spectrotemporal analysis of the recorded waveforms

We visualized the trace recorded from spontaneous sequences produced by a female cuttlefish Sepia officinalis (**Figure S2**A) and the trace produced by a juvenile Sepia bandensis. We computed their spectrograms (**Figure S2A**–**B**). We find that they share energy in the 15–20 Hz range.

## Discussion

In this study, we demonstrated that two species of cuttlefish exhibit comparable stereotyped “arm wave signs” that can be elicited by visual and mechanosensory displays. More specifically, we found that cuttlefish wave back at videos displaying other cuttlefish signing, and that they do so more often in response to the original videos compared with upside-down versions of those videos, and compared with videos depicting other cuttlefish not signing. Furthermore, we found that cuttlefish respond in a similar manner when the signs are conveyed via mechanosensory stimulation in place of visual stimulation. For example, we found that cuttlefish respond more frequently when mechanosensory stimuli are displayed in their original configuration as opposed to a time-reversed version in which traces are played backwards.

The two manipulations of flipping an image upside-down and playing a sound backwards are known to interfere with semantic content, while leaving all other image/sound properties unaffected (79, 80). For example, an upside-down image allocates contrast to detail in a manner identical to that of the upright image, and contains the same set of contours/objects/edges. However, its interpretability is compromised: as convincingly demonstrated by the Thatcher illusion (81), gross distortions of the eye and mouth regions go perceptually unregistered when the face is upside down, but are readily available in the upright configuration. Similar effects can be demonstrated for natural scenes (82, 83). Reverse playback achieves a similar goal ((73, 74)), despite the resulting sound retaining most of the original sound properties such as pitch, loudness, and overall rhythm.

Our finding that cuttlefish respond significantly more to normal versions of visual/mechanosensory stimuli as opposed to their upside-down/time-reversed counterparts (**Figure** 2**D**) indicates that wave signs carry semantic information. In the following, we review relevant connections with the existing literature, address interpretational difficulties, and discuss potential implications of our findings for multimodal communication in cephalopods.

### Relation to previous literature on arm postures/movements in cephalopods

The arm wave signs documented here may represent signals shared across different species of cuttlefish. In addition to the two species tested here, we anecdotally observed a display of similar signs in the cephalopod Sepiola atlantica interacting with a shrimp, see **(Supplementary video 6**). They co-occur with other stereotyped arm wave postures/movements already documented in cephalopods. More generally, arm wave movements and postures for communication are well described and studied in the cephalopod literature. For example, in Sepia officinalis, the “fourth arm extension” (7–9) is well documented and co-occurs often with our fourth signs in males. The “spread of arms and web” belongs to a documented class of reference signs (7) and could potentially overlap with our “Crown” sign category. It could also be similar to the opening and extension of the arm crown in the exaggerated “Umbrella pattern” observed during mating in Sepia latimanus (7). A type of “arm wave” has been documented as posture (8, 9) and may overlap with one of our signs (additional specifications would be needed to ascertain this potential connection). Okamoto and collaborators showed that cuttlefish of the species Sepia pharaonis exhibit a stereotyped and specific arm wave movement during hunting, which the authors interpreted as a mimicking signal (11). In squids, arm postures have been widely studied in the context of communication and camouflage. Mather and collaborators have documented “squid dance” ethograms featuring a “bad hair” display that is to some extent comparable with our “up” wave sign: in this display, the *“arms and tentacles are splayed equally, bent proximally toward the body without systematic spacing, held loosely and twisted”* (15). In the squid Sepiotheutis australis, Jantzen and collaborators have described different postures and movements in the context of mating, potentially constitutive of a complex visual communication system (14). In the squid, Mooney and collaborators have found that a specific frequency of sounds evokes stereotyped fin movements (55) that may share similarities with the movements we evoked during our playback experiment. Our observations are consistent with their findings.

### Candidate structures for mechanoreception of arm wave signs

Our results indicate that arm wave signs can be perceived through mechanoreception, however they do not allow us to pinpoint the physiological mechanism underlying this phenomenon. The two candidate organs for low-frequency (15–35 Hz) sensory perception in cuttlefish are the lateral line and the statocysts (37, 39, 46, 47, 64, 70, 84). The position of the lateral line on the arms and head of the animal constitutes a form-to-function argument in favor of our hypothesis, as does the presence of iridophore cells on the arms, which are used for communication through light polarization (32–34, 36). Furthermore, it is well established that cephalopods possess a wide repertoire of behavioral responses to sounds that could stimulate the lateral line, statocysts, or both (38, 42, 43, 45–47, 49–52).

During the mechanosensory experiments, we observed that cuttlefish contract one or both eyes, suggesting that mechanosensory vibrations may also evoke the vestibulo-ocular reflex via stimulation of the vestibular system (cephalopod statocysts are known to respond to low-frequency vibrations and act as gravity detectors (47, 50)).

Further study and exploration at the cellular and molecular levels should be able to determine whether arm wave signs are perceived through the lateral line, statocysts, or both. Perception of arm wave signs through statocyst stimulation is an equally relevant alternative assumption to lateral line stimulation. For example, there is electrophysiological evidence that statocysts in the Longfin squid can detect low-frequency vibrations (20–1000 Hz) from predators and prey and for navigation (47, 50).

Mooney and collaborators point out that, despite cephalopods not producing sounds, jet propulsion in squid produces low-frequency water flow with strong particle motion that could be sensed and exploited during shoaling (47). Our results are consistent with this notion, particularly in relation to the crown sign, which involves a certain component of jet production. Existing evidence indicates that the working distance of hair cells in the lateral line is one body length, while statocysts are stimulated by whole body displacements at larger working distances and are therefore better suited as receptors for sound (42). In our mechanosensory experiments, the hydrophone/speaker was usually placed at about one body length from the animal, close to the arms. In future experiments, we plan to vary this distance and use the resulting information to disentangle the possible contributions of lateral line versus statocysts for perceiving arm wave signs.

### Potential meaning of arm wave signs

All studies of animal communication suffer from inherent difficulties in interpreting the signs produced by animals, and our study is no exception. With this caveat in mind, we present some hypotheses based on our observations combined with relevant knowledge from the existing literature.

Among various possibilities, the arm wave signs may represent domination displays: we observed that when a cuttlefish is waving, it often places itself in front of the tank while other cuttlefish withdraw, as if the waving animal had established some form of hierarchy. At the same time, cuttlefish of different sizes all produce signs, which is not entirely consistent with the above interpretation. Alternatively, the signs may represent courtship displays, in line with their attractiveness and exaggeration (20, 77). However, we observed that these signs can be expressed by juveniles who have not yet reached sexual maturity in both Sepia bandensis and Sepia officinalis.

In yet another interpretation, the signs could represent aversive displays, as they appear to be expressed towards other cuttlefish in defensive contexts, such as the presence of potential predators (shrimps bigger than juveniles) or humans approaching the tank. It is also possible that some signs may express internal states and or/mood, as they are sometimes spontaneously produced in the apparent absence of external stimulation. Finally, the arm wave signs may be utilized to attract and/or detect prey, as they are often exhibited in the close presence of live prey food (especially we observed juvenile displaying the crown sign as soon as live prey shrimps are inserted in the tank). Furthermore, during both visual and mechanosensory experiments, cuttlefish occasionally elicited other types of arm displays, like hunting arms or tentacule ejection for prey catching. From the above, it is clear that the issue of interpreting the potential meaning of these signs is complex and unlikely to yield a simple answer.

Based on the above considerations, we believe the most plausible interpretation is that these signs carry a variety of possible meanings/functions depending on the associated behavioral contexts. We found that the signs are expressed concomitantly with other body patterns, such as dynamic skin coloration (passing cloud, orange, black spot, dark skin coloration, bright white coloration, zebra bands), changes in body shape (body rounded or flatten) and stereotyped fin flipping. We also observed that they co-occur with other behaviors such as inking (especially during the mechanosensory experiment), pupil contraction, head/eye movements, prey detection, burying, hiding behind rocks, and swimming. These observations align with previous studies on behavioral responses to sound in cuttlefish (54). During both visual and mechanosensory experiments, juvenile Sepia bandensis expressed only the “up” and “crown” signs, despite the fact that they did display the remaining signs spontaneously outside the experiments. Future research will be necessary to explore whether this may be a consequence of their young age, for example, or whether those two signs are functionally different from other signs.

More generally, deciphering wave signs will require careful study and characterization of their association with specific body patterns and behavioral contexts for large datasets, likely through machine learning methods for quantifying dynamic skin patterning (6, 76, 85, 86). Furthermore, specific meaning may be ascribed not just to individual signs, but to combinations of multiple signs: we observed that cuttlefish spontaneously display wave signs in complex long-lasting sequences, the meaning of which may again depend on body patterning and behavioral context (87). Although our results do not speak directly to this possibility and do not support detailed characterization of the rules potentially underlying a putative “syntax” of arm wave signs, the methods presented in this study provide a starting point for developing tools that may enable such a characterization. We expect that this line of inquiry will play a central role in future research aiming to uncover the potential meaning of arm wave signs in cephalopods.

### Convergent evolution of multimodal communication

The nature of the phenomenon exposed by our experiments carries potentially fascinating implications for understanding the evolution of animal communication in wildly diverse systems. The evolutionary and functional connections between hearing and mechanoreception are well established, allowing us to draw a parallel between audiovisual communication in vertebrates and multimodal perception of arm wave signs in cephalopods: in both cases, animals exploit the full range of sensory signals conveyed by communication displays, with a specific focus on visual and mechanosensory signals. Regardless of their evolutionary trajectories, all animals must communicate within the constraints imposed by the physical world. Although these constraints may differ for different animals, they share some fundamental properties that, in turn, dictate the architectural principles underlying sensory systems. Just as the camera-type eye of cephalopods is an example of convergent evolution with the vertebrate eye, we speculate that the lateral line of cephalopods may represent an example of “convergent function” with the vertebrate ear (compared to cephalopod statocysts) or the vertebrate lateral line (compared to cephalopod lateral line), with respect to communication of socially relevant signals. We hope to explore this notion in future research, in particular its potential for facilitating communication between cuttlefish and animal species that may seem immeasurably distant from these creatures.

## Supplementary materials and data availability

Supplementary materials including supplementary videos, a subset of data that allow replication of the results, and coding scripts are available in an open access Mendeley Data repository online: “Cuttlefish interact with multimodal arm wave sign displays”, Mendeley Data, V4, doi: 10.17632/f3sp55762b.4

## Acknowledgments

We thank Chaire Beaut(e)s PSL-L’Oreal for supporting the research documented in this study. We thank German Sumbre and Ehud Vinenpinksy (IBENS) for lending equipment (hydrophone) that allowed us to perform our mechanosensory experiments, and for their assistance in utilizing the equipment. We also thank Jean Michel Maggioranni and his team at the Aquarium de la Rochelle for providing animals and for their help and support with cuttle-fish breeding and husbandry. We thank Balkis Cadi for providing invaluable logistic and administrative assistance and support during the duration of the study.

---

**Fig. S1.**
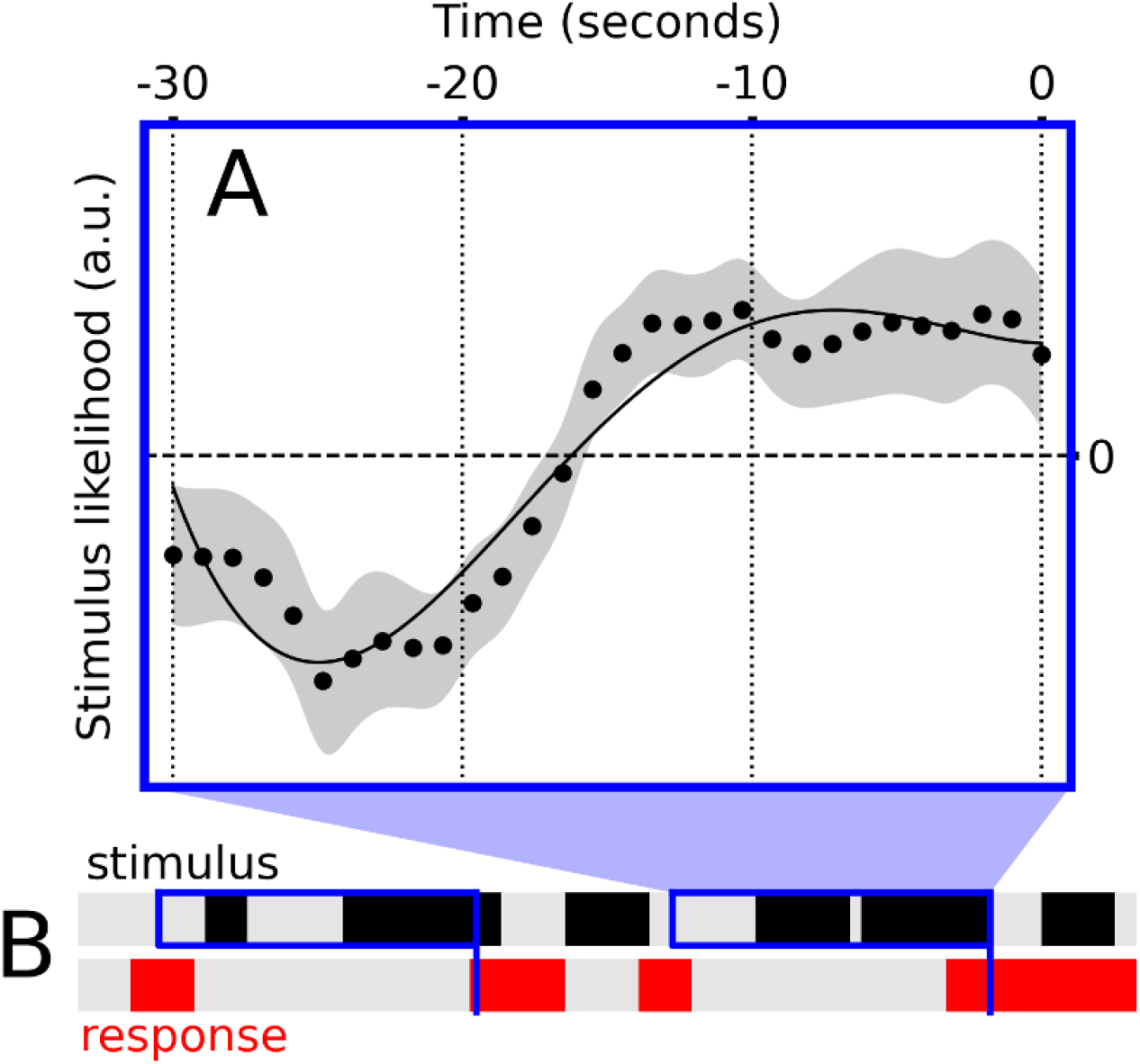
Cuttlefish Sepia officinalis respond to visual wave signing in sluggish fashion over several seconds. We computed the behavioral equivalent of a spike-triggered average: for each occurrence of a wave sign produced by the test animal (indicated by **B**), we extracted the preceding 30-second binary sequence of visual stimulation (absence/presence of wave signing in the visual stimulus, indicated by black segments in **B**). This procedure is indicated by the blue elements in **B**: a time point is selected within the red segments, as indicated by blue vertical lines in **B**, and the preceding 30 seconds of the stimulus are extracted, as indicated by blue rectangles in **B**. We then averaged across all extracted 30-second segments to produce the trace in **A**. This trace was computed separately for each experiment, and we subtracted its mean across time separately for each experiment to normalize its baseline to 0. **A** shows the average across all experiments and animals. We did not measure substantial differences in shape between upright and upside-down traces, so we show the combined case here. Zero on the x axis indicates the production of a wave sign on the part of the test animal. The trace in **A** shows that this response is triggered by stimulation occurring over an extended period of 15 seconds before the event, indicating that test animals do not respond immediately to the sight of a wave sign, but rather integrate the stimulus over several seconds before producing their response. This information was integrated into the analysis that went into **Figure 2D**

**Fig. S2.**
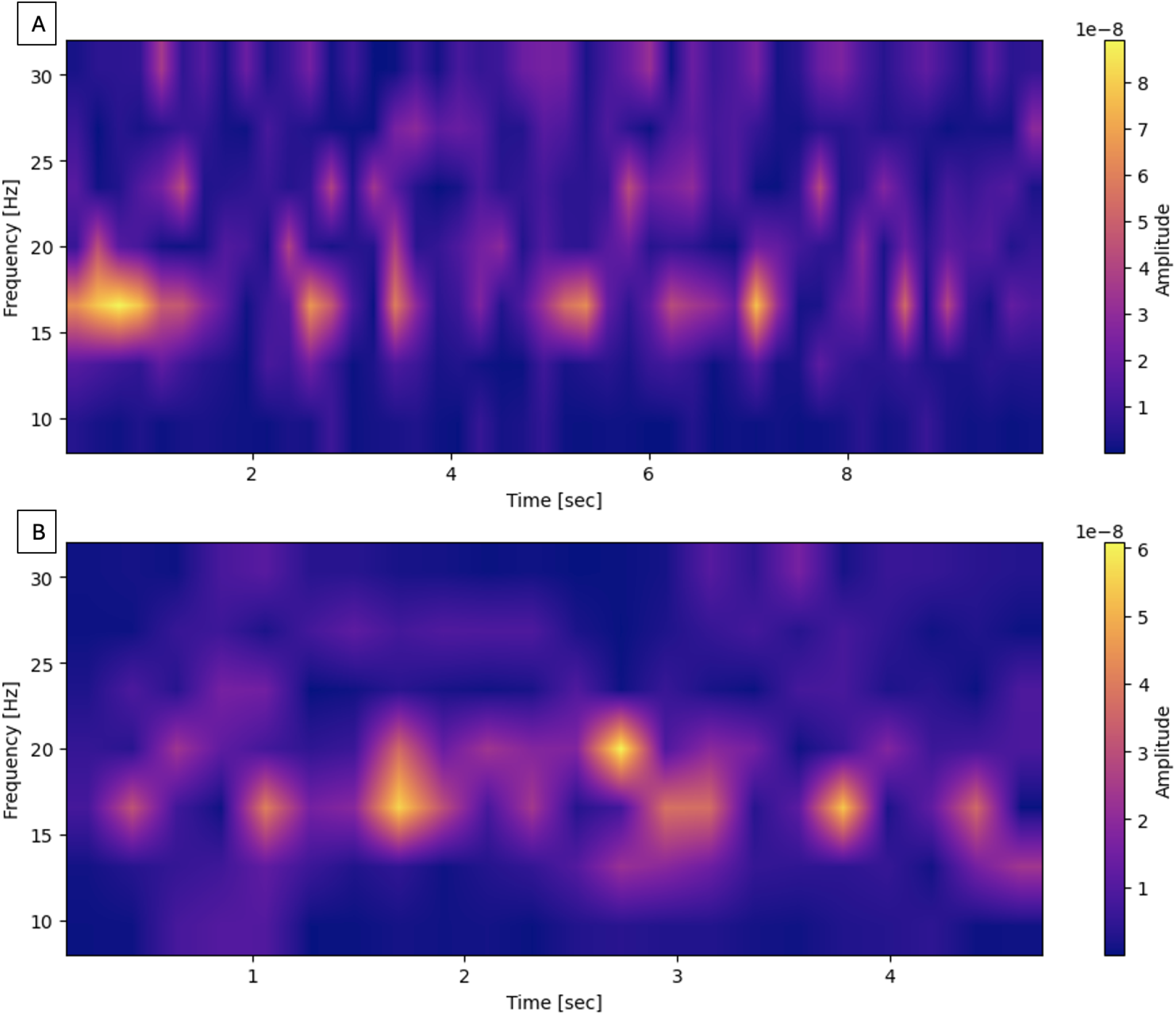
Spectrotemporal content of recorded wave signs. Spectrograms computed from two recorded samples of wave signs (using a hydrophone, see Figure 4A–C) indicate that energy relevant to signing mostly lies within the 10—30 Hz range. Existing literature on the lateral line and statocysts (41, 47, 54) points to this range as the most plausible carrier for information relating to biologically meaningful vibrations like those studied here.

